# Investigating Protein-Protein Allosteric Network using Current-Flow Scheme

**DOI:** 10.1101/259572

**Authors:** Wesley M. Botello-Smith, Yun Luo

## Abstract

Protein dynamic network analysis provides a powerful tool for investigating protein allosteric regulation. We recently developed a current-flow betweenness scheme for protein network analysis and demonstrated that this method, i.e. using current-flow betweenness as edge weights, is more appropriate and more robust for investigating the signal transmission between two predefined protein residues or domains as compared with direct usage of correlation scores as edge weights. Here we seek to expand the current-flow scheme to study allosteric regulations involving protein-protein binding. Specifically, we investigated three gain-of-function mutations located at the binding interface of ALK2 (also known as ACVR1) kinase and its inhibitory protein FKBP12. We first searched for the optimal smoothing function for contact network construction and then calculated the subnetwork between FKBP12 protein and ALK2 ATP binding site using current-flow betweenness. By comparing the networks between the wild-type and three mutants, we have identified statistically significant changes in the protein-protein networks that are common among all three mutants that allosterically shift the kinase towards a catalytically competent configuration.

## Introduction

Among over 500 protein kinases encoded in human kinome, Type 1 Serine/Threonine Kinase Receptors (STKR1) represent an important family that controls the transforming growth factor-β (TGF-β) signaling and bone morphogenetic protein (BMP) signaling. In mammals, STKR1 encompasses seven members named Activin-Like Receptors 1-7 (ALK1-7). The malfunction of these family kinases is often associated with cancer and bone formation disease^1, 2^. A notable case is the genetic gain-of-function mutations (R206H, Q207E, R258G, G328E/W, and G356D) found in ALK2 (also known as ACVR1), which are associated with Fibrodysplasia Ossificans Progressive (FOP), a rare hereditary disorder showing progressive heterotopic ossification in soft tissues, and Diffuse Intrinsic Pontine Glioma (DIPG), a rare childhood brain tumor disorder^3–6 7,8^. Among those mutations, residues 258, 328 and 356 are part of the ATP-binding site, thus alter ALK2 kinase function directly. However, residues 206 and 207 are located on the intracellular glycine-serine rich (GS) domain at the N-terminal of the kinase, away from the ATP-binding site, but close to the binding interface between the kinase and a regulatory protein FKBP12 (a 12-kDa FK506-binding protein).

In the absence of extracellular ligand, STKR1 kinase activity is physiologically inhibited by binding of FKBP12. Extracellular ligand binding facilitates dimerization of type II receptor with STKR1, and subsequent phosphorylation of the GS domain, thus triggering STKR1 activation and downstream signaling. Biochemistry studies have shown that R206H, Q207E mutations in the GS domain do not abolish the FKBP12 binding in absence of type II BMP receptors^4, 9^. Our previous molecular dynamics (MD) simulations and free energy calculations revealed that the conformations of the ATP-binding site of ALK2^R206H^ and ALK2^Q207E^ in presence of FKBP12 are similar as the one of wild-type ALK2 (ALK2^WT^) without FKBP12 bound^10^. More specifically, we found that a salt-bridge R375-D354 between A-loop and the conserved DLG motif is unstable in two mutants. This is an intriguing finding because, despite the fact that R206H and Q207E are located 30 Å away from the A-loop, they are somehow able to bypass FKBP12-mediated inhibition and allosterically shift the ATP-binding site conformation ensemble towards the active state.

Protein dynamical network analysis based on amino acid pairwise correlations or interactions provide a means to understand such allosteric phenomena. Same as in graph theory, a protein network topology is usually defined by a set of nodes, representing certain atoms in amino acids or centers of mass of amino acid side chains, and a set of edges connecting the nodes. Various edges weighting schemes have been previously discussed^11^. To assess the importance of a given node or edge with respect to signal propagation within a protein or between proteins, a common approach is to make use of betweenness centrality, also known as geodesic centrality, which measures the number of shortest paths that crosses a given node or utilizes a given edge in the studied network. However, if one considers only the shortest path between each residue pair, then residues and interactions near to, but not on, those paths would be missed even if they were in fact still relevant. We recently developed a current-flow betweenness scheme, originated from electrical network theory, for protein dynamical network analysis. This method is more appropriate and more robust when only the transmission between a pair of residues or domains is of interest. Using a classic example of an allosteric enzyme imidazole glycerol phosphate synthase (IGPS), we calculated the current-flow betweenness network between an allosteric effector binding site and a remote catalytic site 20 A□ apart and demonstrated a significant improvement in the convergence of the allosteric network topology between independent replicas of simulations^11^.

In current study, we seek to expand the current-flow betweenness approach to a broader range of allosteric regulations involving protein-protein binding. Different from our previous studies^10, 11^, which used a single pair of amino acids as the source and sink for current-flow betweenness calculation, here we report the usage of the current-flow scheme to multiple source/sink pairs in order to study the network change from protein-protein binding interface to the kinase catalytic site. More specifically, we conducted MD simulations of the FKBP12-ALK2^WT^ protein complex, and three gain-of-function mutants at ALK2-FKBP12 binding interface. Using these four systems, we aim to capture the statistically significant network changes that are common among all three mutants with the intent of using this information to reveal the underlying molecular mechanism of the gain-of-function genetic diseases. The three mutants include the aforementioned FOP and DIPG associated disease mutants R206H and Q207E, and an engineered constitutively active mutant Q207D that are extensively used to reproduce the FOP phenotype in mouse model^12^.

To this aim, we first search for the optimal smoothing function for network contact topology construction. Next, the suboptimal paths are calculated between FKBP12 protein and the A-loop R375 residue, which is part of the R-D salt-bridge lock that flips out during kinase activation^10^. Analysis of node and edge usage frequency among these high usage subnetworks shows how a single mutation at the protein-protein binding interface changes the protein-protein communication landscape to allosterically shift the kinase towards catalytic competent configuration. Statistical significance of differences between wild type and mutant networks (*i.e.*, delta-network) was assessed by performing analysis of variance using four blocks of trajectories for each system, followed by significance tests for each individual edge. By doing so, we were able to eliminate the noises in the networks due to thermal fluctuation and to pinpoint the changes in nodes and edges usage that are significant for all three mutants. The python script for calculating current-flow betweenness can be downloaded and used alone (https://github.com/LynaLuo-Lab/network_analysis_scripts/blob/master/python_version) or used together with path-finding program Weighted Implementation of Suboptimal Paths (WISP)^13^ and NetworkView plugin^14^ in VMD program^15^.

## Methods

### System Preparation and Simulation Protocols

The system preparation and MD simulation have been described in our previous work^10^, except for ALK2^Q207D^ system, which is generated here using the same protocol. Briefly, wild-type ALK2 (ALK2^WT^) with FKBP bound structure complex was taken from the crystal structure (PDB ID: 3H9R). The missing A-loop residues 362 to 374 and the β-turn residues 273 to 275 were transplanted from a crystal structure (PDB ID 3Q4U). Each mutant system (R206H, Q207E, and Q207D) was prepared from ALK2^WT^-FKBP using *Molecular Operating Environment* (MOE)^16^ software. 10,000 snapshots from 300 ns MD simulations of were used to determine the protonation state of each amino acid using PROPKA^17, 18^. The p*K*a value the histidine in R206H was previously calculated to be ~6.3 using constant-pH simulation, indicating the dominant protonation state of 206H is the unionized state with hydrogen on the N_δ_ atom^10^. CHARMM-GUI^19^ was used to generate each solvated systems. All simulations employed the all-atom CHARMM C36 force field for proteins and ions, and the CHARMM TIP3P force field ^20^ for water. All MD simulations were performed with NAMD^21^ using periodic boundary conditions under constant temperature and pressure (NPT ensemble). A temperature of 300 K was applied via Langevin thermostat and 1 atm of pressure was maintained via Andersen-Hoover barostat. Long-range electrostatic interactions were treated using the particle-mesh Ewald (PME) method. A smoothing function was applied to both electrostatic and VDW forces over the range of 10 Å to 12 Å. The non-bonded interaction list was updated on every integration step using a cutoff of 13.5 Å. The SHAKE algorithm was applied to all hydrogen atoms. All four systems, wild type and three gain-of-function mutants, were simulated for 300 ns and the last 240 ns was split into four 60 ns windows for statistical analysis.

### Construction of Contact Network Topology

In protein networks, residues within a particular distance of another residue for certain percentage of the time are assumed to influence the communication pathway directly, while residues that do not satisfy these criteria are removed from analysis using a contact map. Therefore, the fluctuation in the contact map has a direct consequence in the instability of the network topology. To build upon the study begun in the prior study of the IGPS system^11^, we investigated the potential utility of an additional layer of contact smoothing to improve the contact map stability.

First, to determine reasonable thresholds for distance smoothing, we computed the minimum residue-residue distances for all residue pair using an average structure of the wild-type extracted from 60-120 ns trajectory. We define the minimum distance between any non-hydrogen atom in one residue of a given residue pair to any non-hydrogen atom in the other residue. A Gaussian kernel density estimate was then constructed (with a .05 Å bandwidth using the *gaussian_kde* function from the python *scipy* library). All the values at inflection points, local minima or maxima are marked in Figure 1 (top). The first peak at 1.4 Å corresponds to the closest contact between neighboring residues, which is the N-C bond length of the peptide bond. The second peak at 2.9 Å likely corresponds to the hydrogen bond distance between residue pairs. Thus the first two peaks in Figure 1 (top) represent common feature in protein structures, thus a cutoff of 4 Å (the local minimum after the 2^nd^ peak at 2.9 Å) would be an appropriate choice of cutoff that is system-independent. The 3^rd^ peak at 7.4 Å and those larger than 7.4 Å are likely reflecting how compact the protein is, and thus depends on the protein shape. In order to investigate the effect of the cutoff distance on the flow-betweenness network, we also chose a larger cutoff of 8 Å (the local minimum after the 3^rd^ peak at 7.4 Å) for comparison. In addition to the two distance cutoff, a linear smoothing function was implemented, which smoothly switch the value between full contact (contact value of 1) at 4 Å or less to no contact (contact value of 0) at 8 Å or greater.

**Figure 1:**
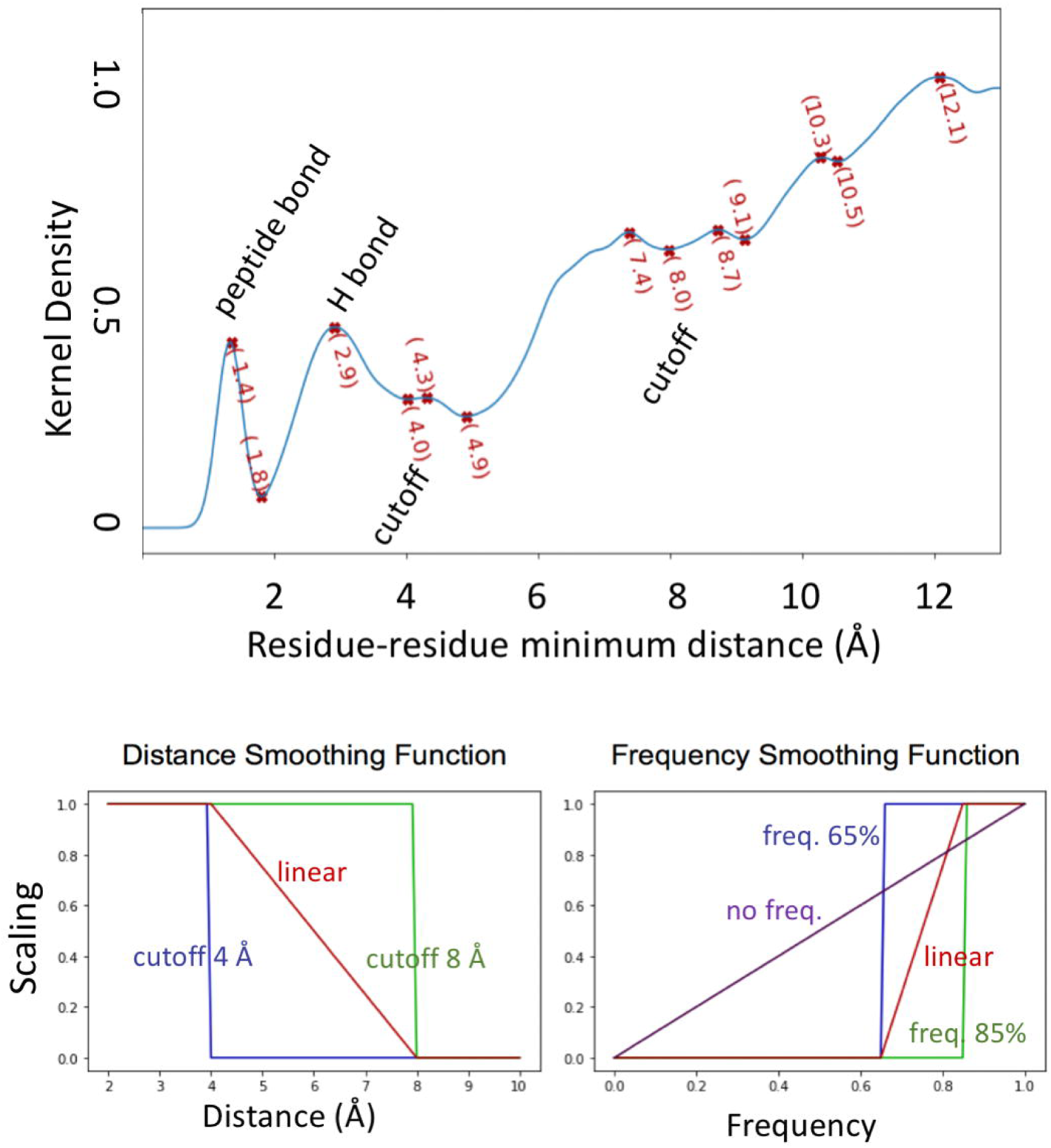
Kernel density estimate of minimal residue-residue distances calculated from an average structure (Top). The graphic representation of the distance (bottom left), and frequency (bottom right) smoothing profiles used in determining network contact topology matrix entries. Here, the ‘linear’ smoothing functions represent interpolating linearly from a value of 1.0 (below the minimum cutoff for distance or above the maximum cutoff for frequency) to a value of 0.0 (above the maximum cutoff for distance or below the minimum cutoff for frequency).

Besides using smoothing function between distance 4-8 Å, we can also use smoothing function on frequencies, for example a pairwise residues within cutoff distance between 65% to 85% of the total time. A linear frequency smoothing scheme was tested which smoothly transitioned from a value of 1 at 85% or greater mean distance contact value to 0 at 65% or lower mean distance contact value (see Figure 1 bottom for a graphical representation). The contact maps for 4 x 60 ns windows of each system would then be constructed by applying the corresponding smoothing scheme.

### Construction of Current-Flow Network

Next, current-flow betweenness for the FKBP12-R375 network was computed using the same protocol as previously described^11^. Briefly, to compute current-flow betweenness, one must first construct the associated adjacency and Laplacian matrices. The adjacency matrix, **A**, is obtained by the resulting matrix of contact values obtained above, multiplied by the pairwise correlation matrix. A diagonal matrix **D** is constructed, with entries equal to the sum of the off diagonal terms of the corresponding rows of **A.** The difference between the diagonal matrix **D** and the adjacency matrix **A** is known as the Laplacian matrix **L.** Next, the inverse of this network Laplacian **L** must be computed to obtain the betweenness of each edge. Here, nodes are defined by α-carbon of amino acids and edges represent the interactions/correlations between nodes. In total, 41 source nodes were defined as all residues of the FKBP protein that contained an atom within 7 Å of any atom of the ALK2 protein in the wild-type average structure. The target node was set to be R375 of the R-D salt-bridge lock in the ALK2 protein. The sub-network of paths leading from FKBP12 to the regulatory salt bridge residue R375 for each system was constructed using the WISP program^13^. The optimal path between each source node on FKBP12 and the sink node R375 is defined by the shortest path, in which the distance is the sum of – *ln*(current-flow betweenness score) for all edges in a given path. A total of 100 paths was then generated by setting the ‘*desired_number_of_paths*’ flag in the WISP input script. We take advantage of WISP program’s fast iterative run of pathfinding routine with gradual increasing pathlength cutoffs until the number of paths found equals or exceeds the given target, and then prunes the paths until only the top 100 paths remain.

### Statistical Analysis of Delta-Networks

In order to assess changes in the allosteric network induced by the three gain-of-function mutations, the mean usage frequency for each edge was calculated over all four windows for each system and was then compared against the corresponding edge in the wild-type. By applying an ANOVA over each individual edge’s values among all windows across all four systems, followed by Tukey’s Honest Significant Differences test, the statistical significance of the change in usage from wild type to mutant was assessed. Any edges with a p-value > 20% (E.g. 80% confidence level) were then filtered out.

The remaining network edges were then exported to a pair of matrix data files, with one file containing the network topology with edge thickness based upon the p-value of each edge, and the other file containing the corresponding difference in edge weights, computed as difference between the wild type value, minus the given mutant value, which we call the “delta-network”. The delta-networks were then visualized using the NetworkView^14^ plugin in VMD program^15^. The p-value matrices served as network topology and edge thickness definitions, while edges were colored according to the difference in either edge value or occurrence frequency accordingly. Similarly, to investigate which nodes exhibited statistically significant differences between wild type and mutant systems, the mean usage frequency of each node was calculated and the analogous ANOVA and Tukey’s HSD tests applied for each individual node.

## Results and Discussion

### Selection of optimal contact topology smoothing using betweenness edge ranking stability

Network constructed based on inter-residue contact maps may exhibit different network topologies as a function of simulation length. Choice of weighting scheme can also impact this phenomenon. Thus, we compare the stability of flow betweenness weights under various distance and frequency smoothing schemes for contact network topology generation. In our previous work^11^, sub-network occurrence frequency was computed in order to provide a consistent edge importance metric to compare correlation and flow betweenness weighting schemes. Here, we directly use the flow betweenness values as a metric for assessing the importance/utility of an edge. Node usage betweenness (*η*) is then calculated as the sum of the betweenness values of all associated edges.

Next, the average node betweenness, <r(*η*)>_s,e_ was computed over all 4 windows for each system, followed by the second round of ranking based on each node’s average *η* ranking R(<r(*η*)>)_s,e_. Finally, the average root mean square deviation of node ranks versus the number of nodes included (Equation 1) was plotted for each system to compare ranking across each contact smoothing scheme. E.g. for each system and smoothing scheme, the top *m* nodes were selected based on the average rank score over all windows and the RMSD of ranking (Rank Error) over four windows was computed using those top m nodes.

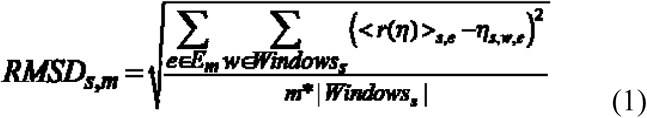

where *s* is one of the systems being considered, *m* is the number of top-ranked edges (by ranking of average *η* values, R(<r(*η*)>)*s,e*, over all windows for system, *s*, being considered), *E*_*m*_, is the set of those top *m* edges for system *s*, and |*Windows*_*s*_| is the number of windows for the system *s*.

Figure 2 shows how the choice of contact network topology construction scheme affects ranking performance using current-flow betweenness weights. Using a 4 Å distance cutoff along with the linear scaling between 65% to 85% frequency yields the lowest Rank Error for all tested systems (highlighted by a yellow box in Figure 2). In essence, this figure illustrates the degree to which ranking of the top N residues varied among the four trajectories for each simulated system. It should be noted that this is by no means an exhaustive set of parameter choices. The choice of distance and frequency cutoffs is still somewhat arbitrary. Future work would require sampling of many different protein systems to ensure an extensive and rigorous examination of optimal smoothing function and distance/frequency cutoff hyperparameters.

**Figure 2:**
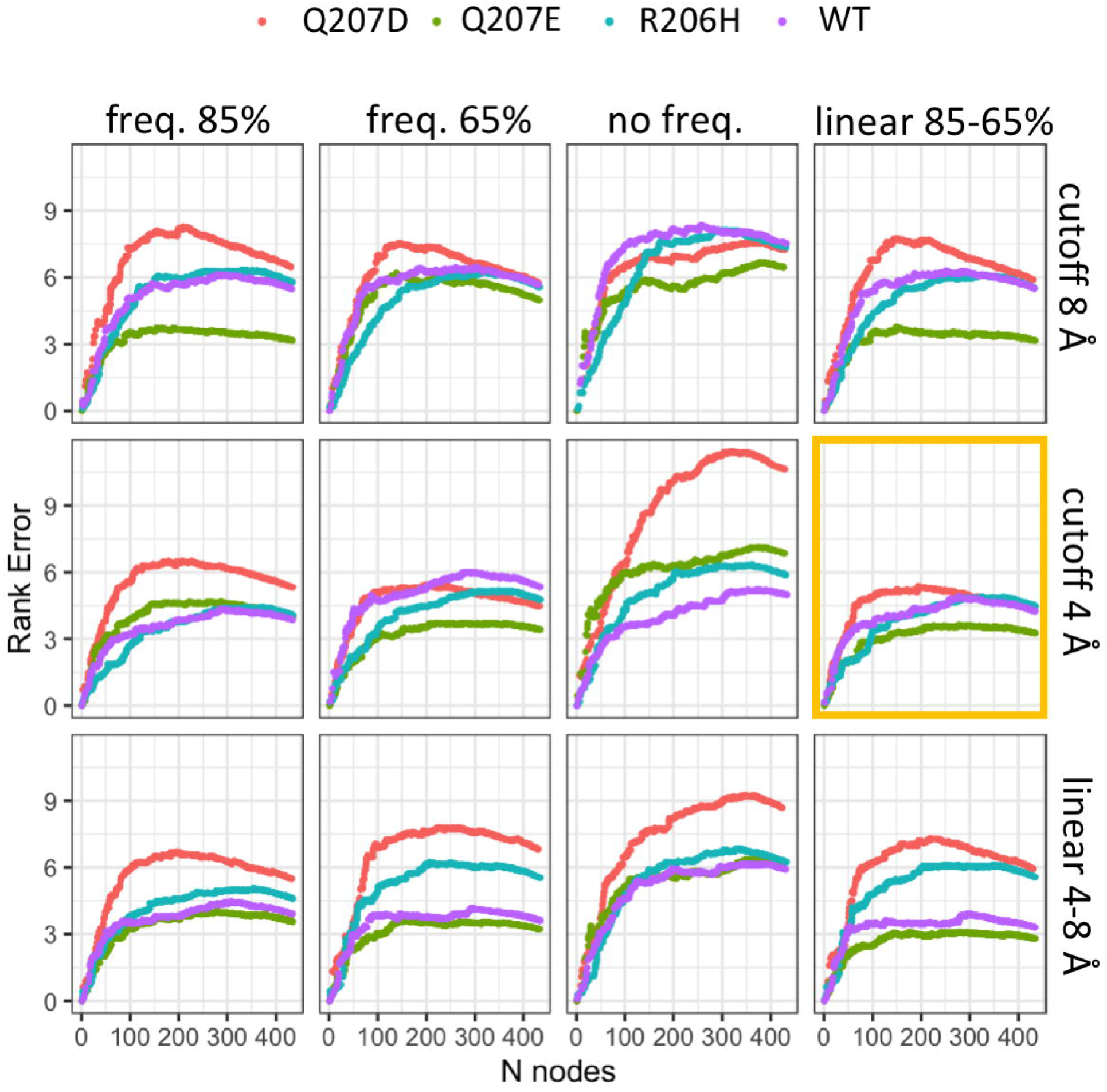
Comparison of node ranking stability under various contact network topology construction schemes (see Fig.1). Subpanel columns indicate frequency smoothing method, rows indicate contact distance smoothing method.

### Comparison of significant change in node usage

Previously, we have shown that R206H mutation decreases the stability of the R375-D354 inhibitory salt-bridge similar as the effect of dissociation of FKBP12 from ALK2. Here we ask whether all three gain-of-function mutations allosterically disrupt the interaction pathway(s) between FKBP12 and A-loop R375 through a similar allosteric network. It was, therefore, expected that FKBP12-R375 interaction pathways generated for these mutants should exhibit significant differences from the wild type. Thus, the subnetwork from FKBP12 protein to R375 is generated for both wild type and mutant. We then subtract the two networks to obtain the difference between wild type and mutant, which we call the “delta-network”.

Changes in node usage are the most readily addressed since there is fewer number of nodes than the number of edges. Figure 3 shows the one-dimensional arrow plot of the significant node usage changes among top 100 paths for each system (left), along with the projection of all nodes for a significant change was observed in at least two of the three mutants (right). Between R206H and Q207E, a few common changes in node usage are observed. There are four gains of usage, occurring at K243 (α helix), N373 (A-loop), and L357 (catalytic loop) in ALK2 and I92 in FKBP12. There are also 3 losses of usage occurring at D241 (α helix) and N372 (A-loop) in ALK2. The node usage changes in Q207D are quite different from the other two systems, with only Y83 in FKBP12 exhibited a loss of usage among all three mutants.

**Figure 3:**
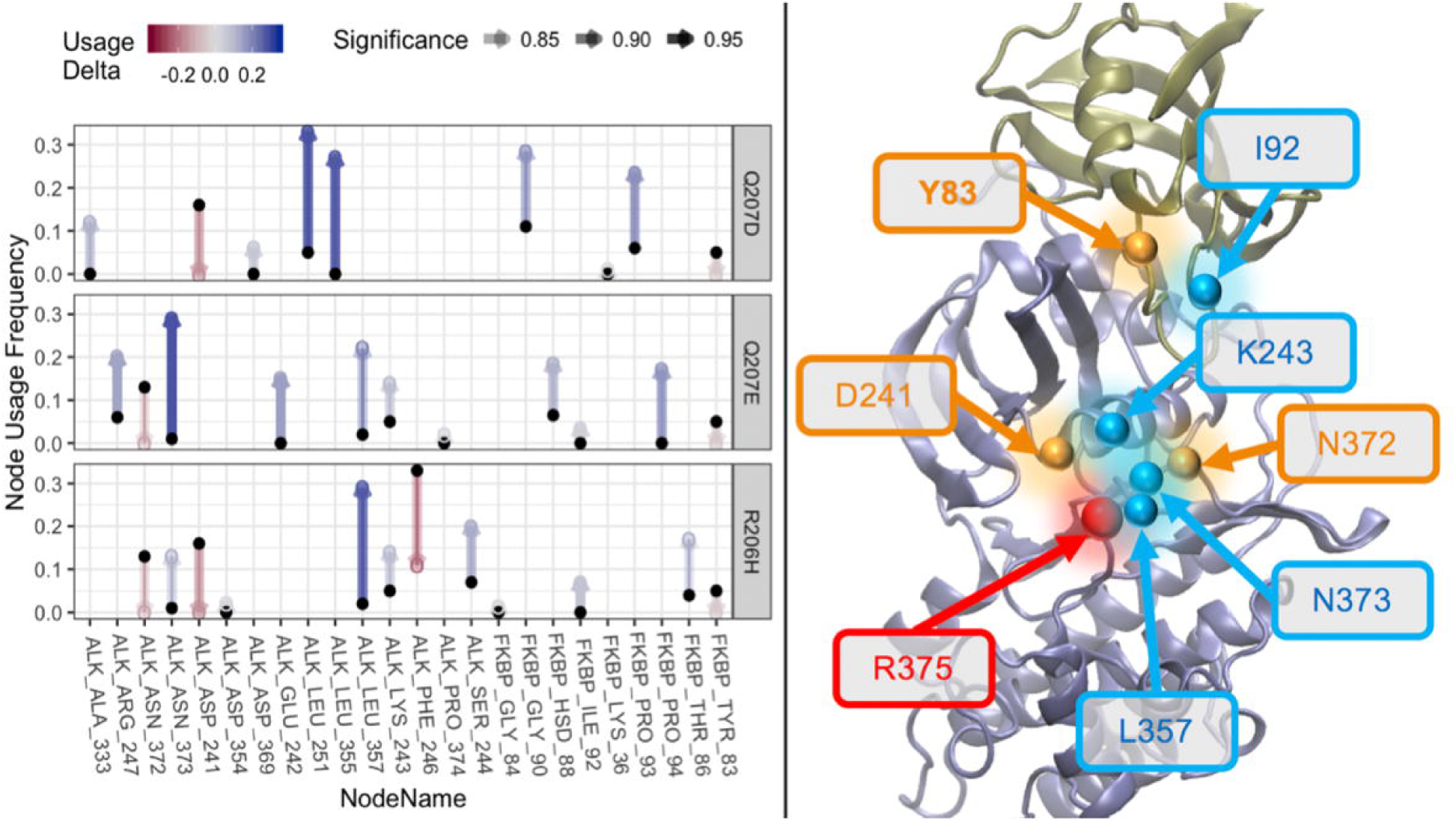
Comparison of changes in FKBP12-R375 node usage among top 100 paths. Arrow direction and color depict the change in the node usage between wild type and the associated mutant. Upward blue arrows represent a gain of usage in mutant and downward red arrows indicate a loss of usage in each mutant. Arrow transparency depicts significance level ranging from a minimum 80% cutoff (full transparency) to maximum 100% (fully opaque).

### Comparison of significant change in edge usage

Next, we investigate whether there are common changes in edge usage among mutants. Comparison of edge usage is somewhat more complicated. This is because there are many more edges than nodes. Edge usage is also much more sensitive to changes in network topology and weighting scheme. For a system of N nodes, there are Nc2 (N choose 2) combinations of node pairs possible in undirected networks, such as the ones discussed here (N^2^ in the case of directed networks). Next, node usage frequency will be equal to one half the sum of connected edge usage frequencies for all non-terminal nodes and equal to the full sum for path terminal nodes. This means that the usages of edges connected to a node may have pronounced changes, yet the node usage may change little or not at all if the signs of those edge usage changes cancel. A good example is the IGPS system^11^. Here, an 80% cutoff was used in gauging statistical significance. This owes to the relatively small number of windows available for each system which limited the power of our statistical tests. While the number of nodes attained even at an 80% cutoff was relatively small, there were significantly more edges exhibiting significant usage changes. Figure 4 shows the edge matrix of each system for which a statistically significant change in usage was observed. The data is provided in table form in **Supporting Information**.

**Figure 4:**
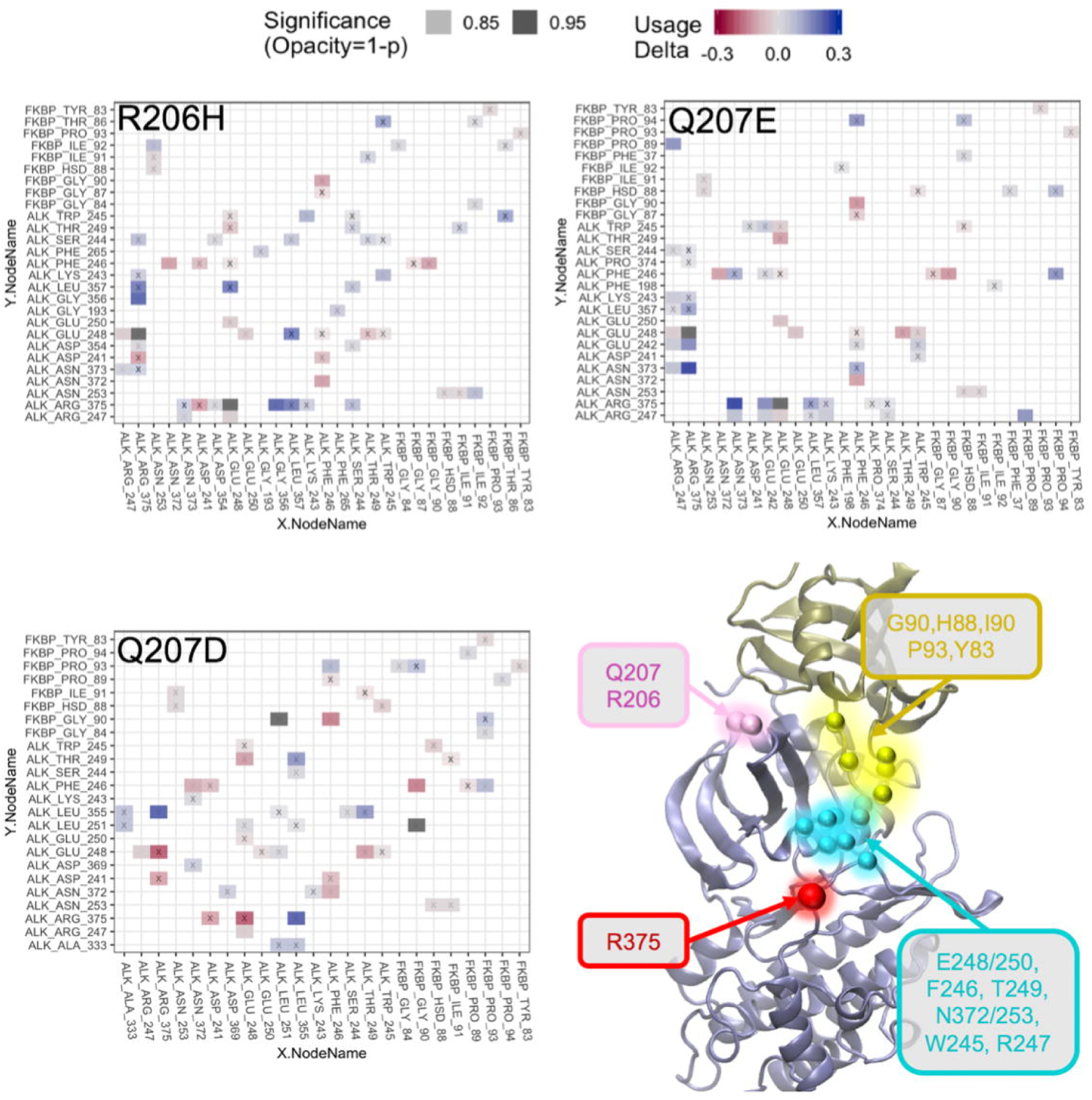
Matrix view of edge usage changes in FKBP12-R375 networks. Transparency indicates relative significance level with 80% significance fully transparent and 100% significance fully opaque. Edges for which significance of change fell below the 80% cutoff have been omitted. Small black ‘x’ symbols have been placed beneath each tile to help visually determine transparency. The color indicates a change in edge usage among top 100 paths between mutant and wild type with blue indicating gain of usage and red indicating loss of usage. The common nodes associated with the edges listed are shown in the bottom right panel. Yellow nodes are the amino acids in FKBP12 and cyan nodes are in ALK2. The source node R375 and two mutation node 206 and 207 are also shown here.

The choice of 80% significance level yielded a rather large number of edges (64 unique edges among all systems), so we limit our discussion to the edges for which a significant change in usage was observed in all mutants. After applying the constraint of edges exhibiting significant changes over all mutant systems studied here, only 10 distinct edges out of 64 were attained (Table 1). Interestingly, all common significant changes for edges usage were negative in sign, indicating a loss of interaction induced by all three gain-of-function mutants. Changes in edge usage appear much more consistent than changes in node usage when comparing all three gain-of-function mutants *vs.* wild type. This observation is in agreement with our previous statement that the edge usage is a more reliable metric than the node usage for comparing changes or perturbations in allosteric network topology^11^.

**Table 1:** Edges exhibiting significant usage changes across all mutants. The residues in the table are highlighted in yellow on the protein structure.

|  |  | Q207D | Q207E | R206H | Mean |
| --- | --- | --- | --- | --- | --- |
| ALK_ARG_375 | ALK_GLU_248 | -0.24 | -0.41 | -0.41 | -0.35 |
| FKBP_GLY_90 | ALK_PHE_246 | -0.15 | -0.13 | -0.12 | -0.13 |
| ALK_THR_249 | ALK_GLU_248 | -0.12 | -0.12 | -0.09 | -0.11 |
| ALK_ASN_372 | ALK_PHE_246 | -0.1 | -0.11 | -0.11 | -0.11 |
| ALK_GLU_250 | ALK_GLU_248 | -0.03 | -0.05 | -0.04 | -0.04 |
| ALK_GLU_248 | ALK_TRP_245 | -0.03 | -0.04 | -0.03 | -0.03 |
| FKBP_PRO_93 | FKBP_TYR_83 | -0.03 | -0.03 | -0.03 | -0.03 |
| ALK_GLU_248 | ALK_ARG_247 | -0.03 | -0.03 | -0.03 | -0.03 |
| FKBP_HSD_88 | ALK_ASN_253 | -0.02 | -0.02 | -0.02 | -0.02 |
| FKBP_ILE_91 | ALK_ASN_253 | -0.02 | -0.02 | -0.02 | -0.02 |

When we plot the nodes associated with those 10 edges in Table 1 on to the 3D structure of protein backbone (Table 1 right), it was quite interesting to see that, although the mutation residues 206 or 207 are located on the left side of the protein, all the edges that have reduced usages induced by mutants are located on the right side of the protein containing the path connecting FKBP12 loop to α helix to A-loop. This observation is in line with our previous work showing that R206H mutation allosterically induces a separation motion between α helix to A-loop^10^. First, the interaction between the sink node R375 on A-loop and E248 on α helix is quite strong in wild type and is significantly diminished in all three mutants. Second, and perhaps more interesting is the disruption of the interaction between G90 on FKBP12 and F246 on α helix. This interaction between FKBP12 loop and α helix has been previously shown to act as a steric barrier to motion of the A-loop which contains R375 of the R-D lock. Also of note is the fourth entry of Table 1 which shows a loss of interaction between residue F246 of the α helix and residue N372 of the A-loop. Therefore current work consolidated this underlying molecular mechanism in which all three gain-of-function mutations disrupt the inhibitory effect of FKBP12 binding protein on the conformation of α helix and A-loop, and thus shift the ATP binding domain towards active conformation.

To illustrate the location of those edges in the edge matrix for each system, we projected the edges onto the three-dimensional protein structures (Figure 5). Nodes exhibiting significant changes in usage among top 100 paths shown as orange spheres. Edge thickness denotes significance level (from 85% to 100%) and edge color denotes the relative magnitude of edge flow betweenness with red representing high betweenness and blue representing low betweenness. On the right side is the delta networks obtained by subtracting the wild type edge betweenness from each mutant, with edge color blue denoting gain of usage in mutant and red representing the loss of usage in each mutant. In delta networks, we see clearly that the edge R375-E248 shows the largest magnitude of the loss of usage among all three mutants. It is worth noting that R375 (A-loop) and E248 (α helix) do not form a direct salt-bridge, but they are part of the ATP-binding site salt-bridge network highly conserved among the STKR1 kinase family^10^. Thus this important change may not be obvious if we only investigate the salt-bridge network. The loss of the indirect coupling between R375 and E248 is likely due to the movement of α helix that was only allowed in gain-of-function mutants.

**Figure 5:**
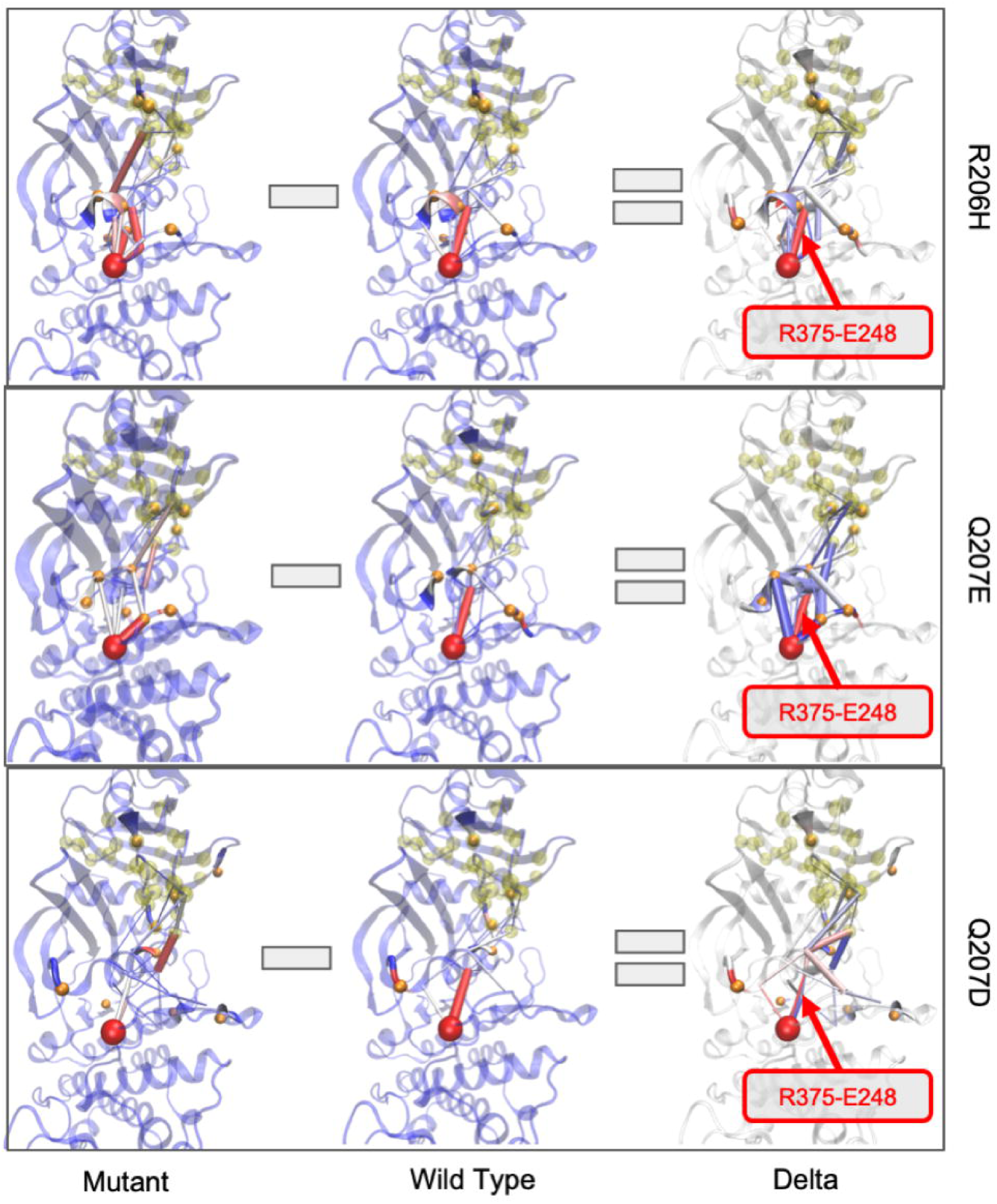
Comparison of FKBP12-R375 delta-networks of top 100 paths for wild type versus R206H, Q207E, and Q207D gain-of-function mutants. **Left and Middle:** Sub-networks of each mutant and wild type system generated using current flow betweenness weights. Source nodes (FKBP12 residues within 7 Å of ALK2) shown in transparent yellow spheres. Nodes exhibiting significant changes in usage among top 100 paths shown as orange spheres. R375 shown as a large red sphere. For mutant and wild type networks, edge color denotes the relative magnitude of edge flow betweenness with red representing high betweenness and blue representing low betweenness. **Right:** In delta networks, edge thickness denotes significance level (from 85% to 100%) while edge color denotes the change in flow betweenness with blue denoting gain of usage in mutant and red representing the loss of usage in the mutant. Edges for which significance of the change in mean usage was below 85% are omitted for clarity. The edge R375-E248 that shows the largest magnitude of loss of usage among all three mutants is indicated on each delta network.

## Conclusion

Network analysis provides a powerful set of tools for investigating dynamical networks of signal propagation in proteins. It was shown previously that the commonly utilized methods for network construction based upon inter-residue contact topology and pairwise correlated motion-based weighting schemes can suffer from instabilities when used in assessing the relative importance of residues and residue to residue interactions (e.g. ranking). Building off our previous work, which made use of a novel current flow betweenness metric to compensate for this instability, we have investigated contact topology smoothing schemes and selected an optimal fit for the ALK2 system based upon node betweenness ranking. This optimized method was then applied to simulations of the wild type FKBP12 bound ALK2 along with three gain-of-function mutant variants to investigate how such mutations disrupt the allosteric regulation of ALK2.

A previous study has shown that the stability R375-D354 salt-bridge at ALK2 ATP-binding site played an important role in the regulation of ALK2 activity. This salt-bridge is normally intact while FKBP12 is bound in wild type, but exhibits marked instability in mutant forms, even in the presence of FKBP12. By employing current flow betweenness, we were able to map the allosteric interaction network connecting residues at the FKBP12-ALK2 binding interface to residue R375 in terms of subnetworks. By performing significance test across four kinase systems and four trajectories in each system, we identified strikingly consistent changes in allosteric network induced by all three gain-of-function mutations. Interesting, although all three mutations are located at the protein-protein binding interface, the perturbed network is on the other side of the protein, containing the path connecting FKBP12 to kinase α helix, and then to A-loop. This information may be useful for future experimental and computational studies that aimed at further elucidating the mechanisms involved in regulation of ALK2 and eventually, development of treatments for those suffering from debilitating mutations of this important signaling protein.

The workflow developed in this study showcased how our current-flow betweenness approach can be used to study a broad range of allosteric regulations within a protein and also between binding proteins. In the latter case, all the amino acids at the protein-protein binding interface can be considered as source nodes that propagating dynamical signals towards a remote active site or ligand-binding site. We also demonstrated here the usage of significance test to capture the network perturbation that is common amongst several mutants vs. wild-type, which is useful to unravel the underlying molecular mechanism of diseases. The method is also sensitive enough to capture the subtle difference in the protein network perturbation between mutants that could be further tested experimentally.

## Supporting Information

A data table containing the significant changes in edge usage between wild type and each mutant. 80% cutoff was used to determine statistical significance. The data is used to plot the edge matrix in Figure 4.

## Acknowledgment

This work was supported by NSF XSEDE research allocation MCB160119.

